# Without quality presence-absence data, discrimination metrics such as TSS can be misleading measures of model performance

**DOI:** 10.1101/235770

**Authors:** Boris Leroy, Robin Delsol, Bernard Hugueny, Christine N. Meynard, Chéïma Barhoumi, Morgane Barbet-Massin, Céline Bellard

## Abstract

The discriminating capacity (i.e., ability to correctly classify presences and absences) of species distribution models (SDMs) is commonly evaluated with metrics such as the Area Under the Receiving Operating Characteristic Curve, the Kappa statistic and the True Skill Statistic (TSS). AUC and Kappa have been repeatedly criticised, but the TSS has fared relatively well since its introduction, mainly because it has been considered as independent of prevalence. In addition, discrimination metrics have been contested because they should be calculated on presence-absence data, but are often used on presence-only or presence-background data. Here, we investigate the TSS and an alternative set of metrics −similarity indices, also known as F-measures. We first show that even in ideal conditions (i.e., perfectly random presence-absence sampling), TSS can be misleading because of its dependence on prevalence, whereas similarity/F-measures provide adequate estimations of model discrimination capacity. Second, we show that in real-world situations where sample prevalence is different from true species prevalence (i.e., biased sampling or presence-pseudoabsence), no discrimination capacity metric provide adequate estimations of model discrimination capacity, including metrics specifically designed for presence-pseudoabsence. Our conclusions are twofold. First, they unequivocally appeal SDM users to understand the potential shortcomings of discrimination metrics when quality presence-absence data are lacking, and we provide recommendations to obtain such data. Second, in the specific case of virtual species, which are increasingly used to develop and test SDM methodologies, we strongly recommend the use of similarity/F-measures, which were not biased by prevalence, contrary to TSS.

## INTRODUCTION

During the last decades, species distribution models (SDMs) have become one of the most commonly used tools to investigate the effects of global changes on biodiversity. Specifically, SDMs are widely used to explore the potential effects of climate change on the distribution of species of concern (Gallon *et al*. 2014), to anticipate the spread of invasive species (Bellard *et al*. 2013), but also to prioritise sites for biodiversity conservation (Leroy *et al*. 2014). Therefore, conservation managers increasingly rely on SDMs to implement conservation strategies and policies to mitigate the effects of climate change on biodiversity (Guisan *et al*. 2013). There are various methodological choices involved in the application of SDMs (e.g., data type and processing, variables, resolution, algorithms, protocols, global climate models, greenhouse gas emission scenarios), which make them particularly difficult to interpret, compare, and assess. However, evaluation of their predictive accuracy is probably a common step to most SDM studies across methodological and technical choices. This evaluation allows us to quantify model performance in terms of how well predictions match observations, which is a fundamental and objective part of any theoretical, applied or methodological study.

To evaluate model predictive performance, the occurrence dataset is often partitioned into two subsets (one for calibrating models, and one for testing) and predictions are assessed in terms of whether or not they fit observations using various accuracy metrics (Araújo *et al*. 2005), a method called cross-validation. Other approaches include calibrating on the full dataset and testing on an independent dataset, or, when the modelled species is a virtual, *in silico*, species (e.g., for testing methodological aspects), directly comparing the predicted distribution with the known true distribution (Leroy *et al*. 2015). Accuracy metrics can be divided into two groups: discrimination vs. reliability metrics (Pearce *et al*. 2000; Liu *et al*. 2009). Discrimination metrics measure classification rates, i.e. the capacity of SDMs to distinguish correctly between presence and absence sites. Reliability metrics measure whether the predicted probability is an accurate estimate of the likelihood of occurrence of the species at a given site. Here, we focus on the issues of discrimination metrics, since they are often used in the SDM literature to test model robustness; however we stress the importance of evaluating reliability (see Meynard & Kaplan 2012 as well as Liu *et al*. 2009), for example with the Boyce index which is probably the most appropriate reliability metric (Boyce *et al*. 2002; Hirzel *et al*. 2006; Cola *et al*. 2016).

Discrimination metrics rely on the confusion matrix, i.e., a matrix comparing predicted versus observed presences and absences (Table 1). Such discrimination metrics have largely been borrowed from other fields of science, such as medicine and weather forecasting, rather than being specifically developed for SDM studies (Liu *et al*. 2009). Three classification metrics stand out in the SDMs literature: Cohen’s Kappa, the Area Under the receiver operating characteristic curve (AUC), and the True Skill Statistic (TSS). The AUC was introduced in ecology by Fielding & Bell (1997) (2,821 citations on Web of Science in June 2017), but has since repeatedly been criticised (Lobo *et al*. 2008, 2010; Jiménez-Valverde 2012) because its dependence on prevalence (i.e., the proportion of recorded sites where the species is present) makes it frequently misused. Cohen’s Kappa has also been repeatedly criticised for the same reason (McPherson *et al*. 2004; Allouche *et al*. 2006; Lobo *et al*. 2010). TSS (Peirce 1884), on the other hand, has fared relatively well since its introduction by Allouche *et al*. (2006) (719 citations in June 2017), mainly because it had been shown as independent of prevalence. However, this claim has recently been questioned because of a flawed testing design (Somodi *et al*. 2017). More recently, all of these metrics have been contested because they should be calculated on presence-absence data, but are often used on presence-only or presence-background data, i.e. data with no information on locations where species do not occur (Yackulic *et al*. 2013; Jarnevich *et al*. 2015; Somodi *et al*. 2017). In these cases, False Positives (FP) and True Negatives (TN) (Table 1) are unreliable, which led Li & Guo (2013) to propose alternative approaches, specifically designed for presence-background models. They proposed the use of *F*_pb_, a proxy of an *F*-measure (“the weighted harmonic average of precision and recall”, Li & Guo (2013)) based on presence-background data, and *F*_cpb_, a prevalence-calibrated proxy of an *F*-measure based on presence-background data. Despite the apparent relevance of Li & Guo’s (2013) metrics (13 citations as of June 2017), the field is still dominated by metrics that have been repeatedly criticised, such as AUC and Kappa, or more recently TSS (*e.g*., D’Amen *et al*. 2015; Jarnevich *et al*. 2015; Mainali *et al*. 2015).

**Table 1.** Confusion matrix used to calculate discrimination metrics.

| Predicted values |  | Sampled data |  |
| --- | --- | --- | --- |
|  |  | Presence | Absence |
|  | Presence | True Positives (TP) | False Positives (FP) |
|  | Absence | False Negatives (FN) | True Negatives (TN) |

With this forum, our aim is twofold: (1) illustrate with examples and simulations that, contrary to early claims, TSS is in fact dependent on prevalence, and (2) evaluate an alternative set of metrics based on similarity indices, also known as *F*-measures in the binary classification literature, as potential alternative measures of model predictive ability. Similarity indices assess the similarity of observed and predicted distributions, and can be partitioned into two components to evaluate model characteristics: Over Prediction Rate (OPR) and Unpredicted Presence Rate (UPR). We compare the performance of TSS and similarity-derived metrics on three modelling situations corresponding to the most common modelling setups, depending on the interplay between species and sample prevalence (see below). We finally discuss the applicability of these discrimination metrics in SDM studies and provide practical recommendations.

## SPECIES AND SAMPLE PREVALENCE

Here we will define *species prevalence* as the ratio between the species area of occupancy (AOO, i.e., the area within the distribution of a species that is actually occupied) and the total study area (see Rondinini et al. 2006 for definitions). For example, if the study area encompasses Europe and we have divided the study area into 1-km grid cells, and if we are studying a species that occupies only 15% of those grid cells its prevalence would be 0.15. Notice that species prevalence will vary depending on the resolution of the gridded data and on the extent of the study area. In practice, however, species prevalence is never known, because the true AOO is generally not known, except for the specific case of virtual species (Leroy *et al*. 2015). Hence, for real species, only the *sample prevalence* is known, which is the proportion of sampled sites in which the species has been recorded. Meynard and Kaplan (2012) showed with virtual species that sample prevalence should be similar to species prevalence to produce accurate predictions. However, in practice, we expect sample prevalence to be different from species prevalence, unless the sampling of presences and absences is perfectly random throughout the entire study area. Indeed, samplings of species presences are generally spatially biased (Phillips *et al*. 2009; Varela *et al*. 2014). For example, ecologists look for their species of interest in sites where they have a sense a priori that they will find it, which will inevitably result in a mismatch between sample and species prevalence. Furthermore, a substantial proportion of SDM studies rely on presence-only modelling techniques, which requires to sample ‘pseudo-absence’ or ‘background’ points (hereafter called pseudo-absences). In such cases the sample prevalence is artificially defined by the number of chosen pseudo-absences, and is thus unlikely to be equal to species prevalence.

Neither species prevalence nor sample prevalence should influence accuracy metrics. In the following, we investigate three different cases corresponding to the most common situations of SDM evaluation. First, we investigate the ideal ‘presence-absence’ case where species prevalence is equal to sample prevalence; this case corresponds to well-designed presence-absence samplings or to the evaluation of SDMs based on virtual species where the true AOO is known. Second, we investigate ‘presence-absence’ situations where sample prevalence differs from species prevalence. Last, we investigate ‘presence only’ situations where sample prevalence differs from species prevalence.

## PRESENCE-ABSENCE, SPECIES PREVALENCE = SAMPLE PREVALENCE

In this first case, we define the sample confusion matrix as perfectly proportional or equal to the true confusion matrix, i.e. the entire predicted species distribution is compared to the true species distribution. In practice, this case occurs when the sampling is perfectly random throughout the landscape and species detectability is equal to one, or when evaluating SDM performance with virtual species (e.g., Qiao et al., 2015). With this first case we can analyse the sensitivity of discrimination metrics to species prevalence only.

### The unexpected dependence of TSS on prevalence

Previous studies have already shown that common discrimination metrics such as Kappa and AUC are influenced by species prevalence (e.g., Lobo *et al*. 2008, 2010). However, TSS has been widely advocated as a suitable discrimination metric that is independent of prevalence (Allouche *et al*. 2006). Here we demonstrate with simple examples that TSS is itself also dependent on species prevalence. When species prevalence is very low (and so is sample prevalence), we expect the number of True Negatives (Table 1) to be disproportionately high. In these cases, specificity will tend towards one, and TSS values will be approximately equal to sensitivity (Table 2). As a result, TSS values can be high even for models that strongly overpredict distributions. Figure 1 represents graphically some examples of how overprediction and underprediction play into TSS performance. For example, Fig. 1a shows a model that strongly overpredicts the distribution, producing 300% more False Positives than True Positives, and yet TSS is close to 1 (Fig. 1a, TSS=0.97). Such a high value can in turn be produced by a model which correctly predicts the true distribution with few overpredictions (Fig. 1b, TSS = 1.00). In addition, the over-predicting model (Fig. 1a) will also have higher TSS values compared to a model that only missed 15% of presences (Fig. 1c, TSS=0.85). Furthermore, for identically-performing models, if sample prevalence decreases (from 0.25 to 0.01), then the proportion of True Negatives is increased, and consequently TSS values increased from 0.60 to 0.70 (Fig. 1d-f). Consequently, TSS values can be artificially increased by decreasing sample prevalence. As an unexpected consequence, for two species with different AOO in the study area (thus different sample prevalence), the species with the smaller distribution will be considered better predicted than the one with a larger distribution (Fig. 1d-f).

**Table 2.** Existing discrimination metrics. TP = True Positives, FN = False Negatives, FP = False Positives, TN = True Negatives, *P* = number of sampled presences, *A* = number of sampled pseudoabsences, *prev*_sp_ = estimate of species prevalence.

| Metric | Calculation | References |
| --- | --- | --- |
| Sensitivity | $Sn = TP / (TP+FN)$ | Fielding & Bell (1997) |
| Specificity | $Sp = TN / (TN+FP)$ | Fielding & Bell (1997) |
| True Skill Statistic | $TSS = Sn + Sp - 1$ | Peirce (1884), Allouche *et al.* (2006) |
| Jaccard's similarity index | $Jaccard = TP / (FN+TP+FP)$ | Jaccard (1908) |
| Sørensen's similarity index,<br>$F$-measure | $Sørensen = 2TP / (FN + 2TP + FP)$ | Sørensen (1948), Li & Guo (2013) |
| Proxy of $F$ measure based on presence-background data | $F_{pb} = 2 \times Jaccard$<br>$F_{cpb} = 2 \times TP / (FN + TP + c \times FP)$<br>where $c = P / (prev_{sp} \times A)$ | Li & Guo (2013) |
| Overprediction Rate | $OPR = FP / (TP+FP)$ | Barbosa *et al.* (2013) |
| Underprediction Rate | $UPR = FN / (TP+FN) = 1 - Sn$ | False Negative Rate in Fielding & Bell (1997) |

**Figure 1.**
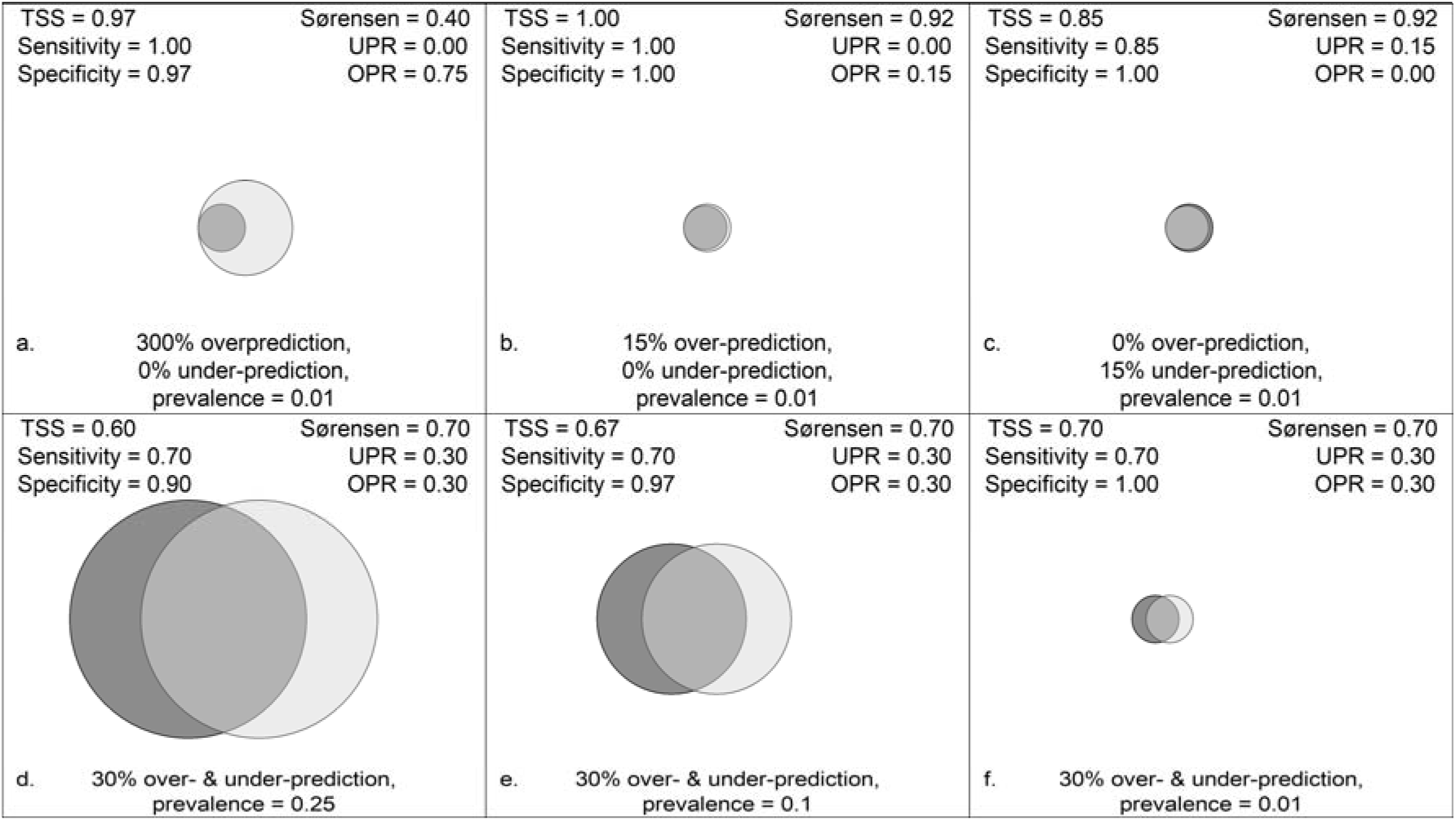
Examples of model performances and associated metrics. The dark grey filled circle represents the proportion of actual presences in the sample. The light grey filled circle represents the proportion of predicted presences in the sample. Therefore, the overlap between the two circles represents the proportion of actual presences correctly predicted as presences (‘True Positives’), whereas the white area represents the proportion of actual absences correctly predicted as absences (‘True Negatives’). At low prevalence (0.10), TSS does not penalise overprediction: a model that strongly overpredicts distribution (Fig. 1a; 300% more False Positive than True Positives) can have a very high TSS (0.97), which is almost equivalent to a model with little overprediction (Fig. 1b, TSS = 1.00). TSS does penalise underprediction (Fig. 1c, TSS = 0.85) much more than overprediction (Fig. 1a-b). For identically-performing models (i.e., similar rates of over- and underprediction), if prevalence decreases (from 0.25 to 0.01) with increasing numbers of True Negatives, TSS values increased from 0.60 to 0.70 (Fig. 1d-f). In other words, for two species with different AOO in a given study area, the species with the smaller distribution have a higher TSS than the one with a larger distribution. Sørensen, on the other hand, accurately discriminates between highly over-predicting and well performing models (Fig. 1a-c). Similarity indices penalise identically over- and underprediction (Fig. 1b-c). In addition, when species prevalence is artificially increased for identical models, both indices remain identical (Fig. 1d-f).

To summarise, TSS values can be misleading in situations where the number of True Negatives is high by (i) not penalising overprediction and (ii) assigning higher values to species with smaller prevalence for identical discrimination accuracy. These flaws can be strongly problematic for ecologists, and during SDM performance evaluation it is generally preferable to assume that overprediction should be equivalent to underprediction (e.g., Lawson et al., 2014). Therefore, we conclude that TSS is prone to similar shortcomings as AUC and Kappa when it comes to its dependence on sample prevalence and AOO.

### Similarity metrics as an alternative

To avoid these shortcomings, we propose to focus the evaluation metrics on three components of the confusion matrix (Table 1): True Positives, False Positives and False Negatives, neglecting the True Negatives that could be easily inflated. In particular, we seek to maximise True Positives, and minimise both False Positives and False Negatives with respect to True Positives. This definition exactly matches the definition of similarity indices from community ecology, such as Jaccard and Sørensen indices or the *F*-measure indices (Table 2). This definition also matches the indices identified by Li & Guo (2013) as potential presence-background metrics. The *F*_pb_ index is in fact equal to twice the Jaccard index (eqn. 13 in Li & Guo 2013), while the *F* index is equal to the Sørensen index of similarity (eqn. 4 in Li & Guo 2013) (Table 2).

Similarity indices have two main benefits. First, their conceptual basis is easy to understand: they measure the similarity between predictions and observations. A value of 1 means predictions perfectly match observations, without any False Positive or False Negative. A value of 0 means that none of the predictions matched any observation. The lower the similarity value, the higher the number of False Positives and False Negatives, proportionally to the number of True Presences. Second, as they do not include True Negatives, they are not biased by a disproportionate number of True Negatives. In return, they do not estimate the capacity of models to correctly predict absences. To illustrate this, we calculated the Sørensen index of similarity (F-measure) on the same examples as above. Sørensen accurately discriminated between highly over-predicting and well performing models (Fig. 1a-c). In addition, when species prevalence was artificially increased for identical models, both indices remained identical (Fig. 1d-f).

Because the specific objectives of SDM studies can be very different (e.g., invasion monitoring versus habitat identification for threatened species), in a particular context we may be more interested to assess whether predictions tend to over- or underestimate observations. Such additional information can be obtained with similarity metrics by partitioning them into two components: overprediction rate and unpredicted presence rate (Table 2). The overprediction rate measures the percentage of predicted presences corresponding to false presences, and was already recommended for assessing model overprediction (Barbosa *et al*. 2013). The unpredicted presence rate measures the percentage of actual presences not predicted by the model, and is also called the false negative rate (Fielding & Bell 1997). Taken together these metrics provide a full view of model discrimination accuracy and allow interpreting the results in the specific context of the study.

### Demonstration based on simulations

To validate these theoretical demonstrations, we performed simulations of the metrics for three case studies with different performances: a first model with 40% overprediction and 40% underprediction, a second model with 40% underprediction and no overprediction, and a third model with 40% overprediction and no underprediction. The first case addresses a predicted range that is shifted in space with respect to the real one; the second and third cases address situations where the predicted range is, respectively, smaller or larger than the real one. For each model, we predicted the distribution range of theoretical species with different prevalence (from 0.01 to 0.60 with a step of 0.01) over an area of 100 000 pixels. Then, for each species, we randomly sampled 500 presences in the total area and a number of absences verifying the condition that the sample prevalence is equal to species prevalence. We repeated this procedure five times. For each repetition, we calculated the True Skill Statistic and the Sørensen index (R scripts available at https://github.com/Farewe/SDMMetrics).

Our results (Figure 2) showed that TSS values decreased with prevalence for cases that overpredicted species distributions, but not for cases that only underpredicted distributions (Figure 2a). This result confirms our expectation that TSS does not penalise overprediction at low prevalence. Sørensen values, on the other hand, remained similar regardless of species prevalence (Figure 2b). These results confirm that in the ideal situation where species prevalence = sample prevalence, the Sørensen index of similarity is a more appropriate metric of model discrimination capacity.

**Figure 2.**
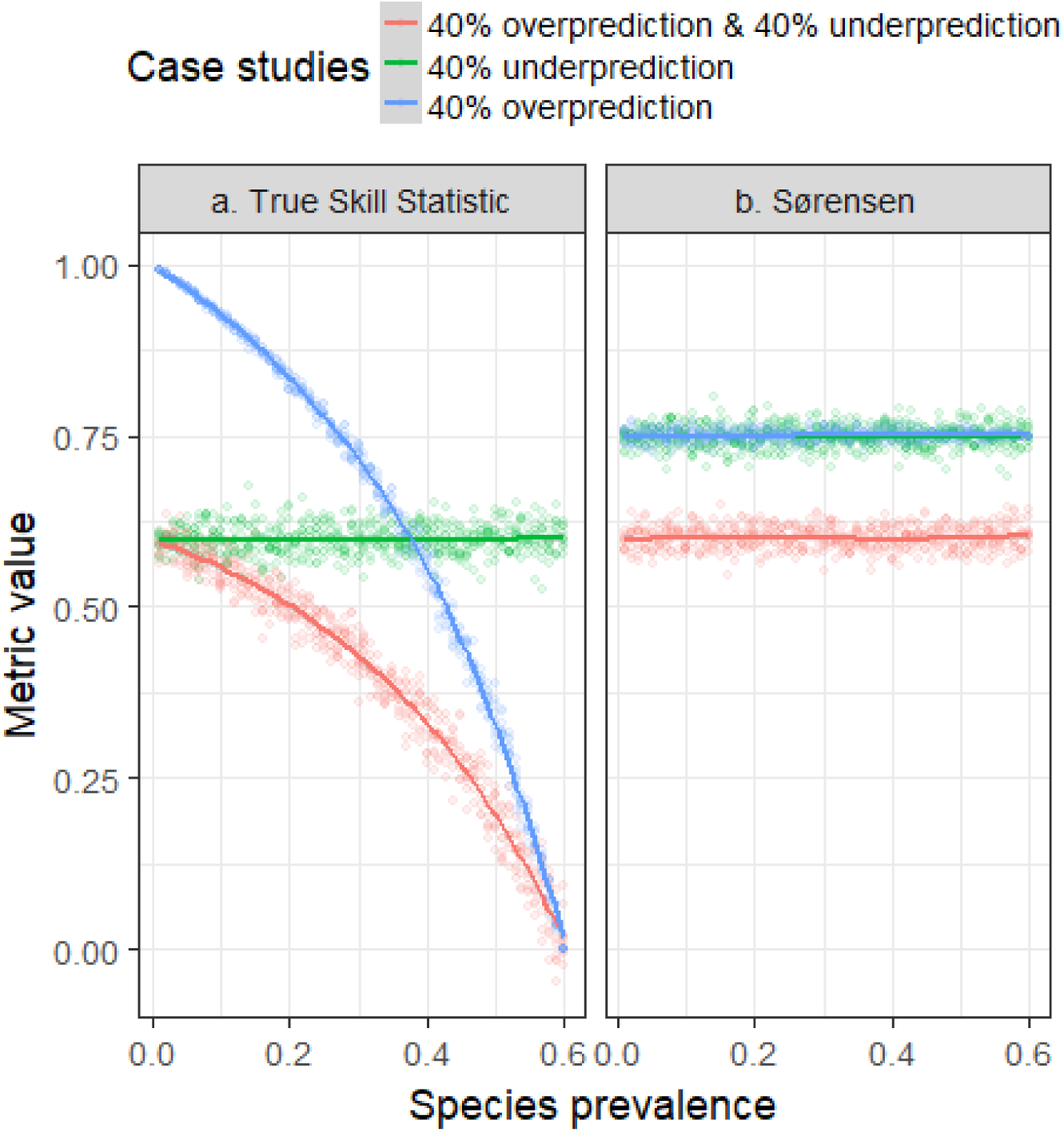
Simulations of the effect of species prevalence on species distribution model discrimination metrics ((a) TSS and (b) Sørensen, equations available in Table 2) in a presence-absence scheme where sample prevalence is equal to species prevalence. Three case studies with varying degrees of over- and underprediction are applied to theoretical species with prevalence ranging from 0.01 to 0.60 with a step of 0.01. The upper limit of 0.60 was chosen so that we can calculate values for models with 40% overprediction. For each species, an evaluation dataset was composed of 500 presences randomly sampled in the total area and a number of randomly sampled absences verifying the condition that the sample prevalence is equal to species prevalence, with 5 repetitions for each species (R scripts available at https://github.com/Farewe/SDMMetrics). These simulations showed that TSS attributes higher values at lower prevalence for case studies that overpredict species distributions, but not for case studies that have only underprediction (Figure 2a). Sørensen values, on the other hand, remain similar regardless of species prevalence (Figure 2b).

## PRESENCE-ABSENCE, SPECIES PREVALENCE ≠ SAMPLE PREVALENCE

When sample prevalence is different from species prevalence, the ratio of sampled absences over sampled presences is different from the ratio of true presences over true absences. For example, if too many absences are sampled (sample prevalence lower than species prevalence), then the numbers of False Positives and True Negatives will be too large compared to True Negatives and False Positives. The major consequence of this mismatch is that any metric comparing sampled presences and absences will not reflect true model performance, unless it contains a correction factor for the mismatch between sample and species prevalence. Note, however, that metrics focusing only on sampled presences (omitting sampled absences) will not be affected by this bias (for example, sensitivity or rate of unpredicted presences will not be affected). We illustrate in Appendix A how the aforementioned metrics are biased by prevalence in this situation: the lower the prevalence, the higher the metric. We further show that an appropriate estimation can only be obtained when an accurate estimation of species prevalence is available, which is generally not the case (see section Estimations of species prevalence).

## PRESENCE-PSEUDOABSENCE OR PRESENCE-BACKGROUND, SPECIES PREVALENCE ≠ SAMPLE PREVALENCE

In presence-pseudoabsence schemes, sample prevalence is highly unlikely to be equal to species prevalence, thus the previous bias also applies in this situation. Furthermore, an additional bias is added by the fact that pseudo-absence points may be actual presence points. This bias will further impact the estimation of False Positive by generating “False False Positives” (FFP), *i.e*. predicted presences corresponding to actual presences but sampled as pseudo-absences. We illustrate with simulation how this bias increases the dependence on prevalence of existing metrics in **Appendix** B, including the prevalence-calibrated *F*_cpb_ metric specifically designed for presence-background (Li & Guo 2013). We also illustrate that a mathematical correction could be applied but requires ideal conditions unlikely to be obtained (perfectly random samplings of presences and pseudoabsences; multiple repetitions; accurate estimation of species prevalence) (see section Estimations of species prevalence).

## ESTIMATIONS OF SPECIES PREVALENCE

The only way to correct discrimination metrics in cases where sample prevalence is different from species prevalence requires an estimate of species prevalence. In presence-absences schemes, species prevalence is usually estimated from the sample of presences and absences – however we assumed here that in many situations this estimate may be biased. Besides, in presence-pseudoabsence schemes this estimation is not available. An alternative approach consists in estimating species prevalence from the modelled species distribution (e.g., Li and Guo, 2013; Liu et al., 2013). Li and Guo (2013) demonstrated that this approach yielded satisfactory results for presence-pseudoabsence based on the *F*_pb_ index. However, these results were later contested by Liu et al. (2016) who found that neither *F*_pb_, nor a TSS-derived metric were able to correctly estimate species prevalence with presence-pseudoabsence data. This inability to estimate species prevalence from presence-pseudoabsence data was expected because an accurate estimation would require strong conditions which are unlikely to be met in reality (see Guillera-Arroita et al., 2015 for a demonstration). Actually, for both presence-pseudoabsence and presence-absence data, estimating species prevalence could be feasible from limited presence-absence surveys, but may be prohibitively difficult or expensive to obtain (Phillips & Elith 2013; Lawson *et al*. 2014). This barrier to estimate species prevalence severely limits the applicability of discrimination metrics for presence-absence and presence-pseudoabsence models where sample prevalence is different from species prevalence.

## DISCUSSION AND RECOMMENDATIONS

In this paper, we have demonstrated that evaluating model discrimination capacity (i.e., the capacity to accurately discriminate between presence and absence) depends on the interplay between sample and species prevalence. We studied three general situations that modellers are expected to encounter in their modelling exercises: (i) a presence-absence scheme where sample prevalence is equal to species prevalence – this situation corresponds to perfectly random presence-absence samplings with no detection bias, or to evaluations based on virtual species; (ii) a presence-absence scheme where sample prevalence is different from species prevalence – a likely situation for presence-absence modelling; and (iii) a presence-pseudoabsence scheme where sample prevalence is different from species prevalence – the general case for presence-pseudoabsence or presence-background modelling.

Our simulations unequivocally indicate that when sample prevalence is different from species prevalence, none of the tested metrics are independent of species prevalence, corroborating previous conclusions on the TSS (Somodi *et al*. 2017), and invalidating the propositions of Li and Guo (2013). Our rationale and conclusions on TSS relate in fact to the same argumentation as provided on AUC by Lobo et al. (2008). Both TSS and AUC have the same shortcomings. Most importantly, Lobo et al. (2008) showed that the total extent to which species are modelled highly influenced AUC values. Indeed, the total study extent drives species prevalence (termed Relative Occurrence Area in Lobo et al. 2008); increasing extent reduces species prevalence and vice versa. Consequently, artificially increasing the modelling extent will artificially decrease prevalence, which in turn will increase AUC values (Lobo *et al*. 2010; Jiménez-Valverde *et al*. 2013), but also TSS values as we showed here. Likewise, comparing species with different AOO over the same extent will provide an unfair advantage to species with smaller AOO because they will have a smaller prevalence. In fact, these shortcomings are likely to be derived to any measurement that need to estimate either FP or TN (Jiménez-Valverde *et al*. 2013).

Our first recommendation is a compelling advocacy for improving data quality in SDMs. Our arguments as well as those of Lobo et al. (2008, 2010) and Jiménez-Valverde et al. (2013) suggest that the quest for an ideal discrimination metric is futile, unless reliable presence-absence data is available. Indeed, an unbiased set of presence and absence data is required to estimate species prevalence (Guillera-Arroita *et al*. 2015), and any metric based on TN and FP (Jiménez-Valverde *et al*. 2013). Therefore, we advocate the importance of collecting more informative data. Ideally, we emphasise the necessity of obtaining at least a random or representative sample of presences and absences (Phillips & Elith 2013), or to improve data collection, for instance, by recording non-detections to estimate sampling bias and species prevalence (Lahoz-Monfort *et al*. 2014; Guillera-Arroita *et al*. 2015). Cross-validation procedures can lead to overoptimistic evaluations because of data autocorrelation, and specific procedures can be applied to avoid this further bias (Roberts *et al*. 2016). We also emphasise the importance of appropriate spatial extent; although a framework to choose spatial extent does not exist, guidelines exist to improve spatial extent definition (Barve *et al*. 2011; Jarnevich *et al*. 2015).

Our second recommendation concerns the case where quality presence-absence data are available. This is also the case of virtual species, which are increasingly used to develop and test SDM methodologies (Li & Guo 2013; Meynard & Kaplan 2013; Varela *et al*. 2014; Miller 2014; Leroy *et al*. 2015; Liu *et al*. 2016; Ranc *et al*. 2016; Hattab *et al*. 2017). Our results unequivocally demonstrated that similarity/F-measure metrics, and their derived components (OPR, UPR) were unbiased by species prevalence and can thus be applied in these cases as discrimination metrics with better results than the classic Kappa, AUC and TSS metrics. Therefore, we strongly recommend the use of these metrics in the specific case of virtual species. After all, virtual species are used to demonstrate the shortcoming and/or advantages of some methods over others, and therefore the use of appropriate evaluation metrics is highly desirable.

